# Cyclodextrin polymer coatings resist protein fouling, mammalian cell adhesion, and bacterial attachment

**DOI:** 10.1101/2020.01.16.909564

**Authors:** Greg D. Learn, Emerson J. Lai, Horst A. von Recum

**Affiliations:** Department of Biomedical Engineering, Case Western Reserve University, Cleveland, OH, USA

**Author notes:** Corresponding Author, 10900 Euclid Avenue, Cleveland, OH 44106.

**Keywords:** Biomaterial, Polymer, Polysaccharide, Surface, Coating, Antifouling

## Abstract

Undesired attachment of proteins, cells/bacteria, and organisms on material surfaces is problematic in industrial and health care settings. In this study, polymer coatings are synthesized from subunits of cyclodextrin, an additive/excipient found in food/pharmaceutical formulations. These unique polymers, which have been applied mainly towards sustained drug delivery applications, are evaluated in this study for their ability to mitigate non-specific protein adsorption, mammalian cell (NIH/3T3) adhesion, and bacterial cell (*Staphylococcus aureus, Escherichia coli*) attachment. Effects of cyclodextrin polymer composition, particularly incorporation of nonpolar crosslinks, on material properties and passive anti-biofouling performance are investigated. Results suggest that lightly-crosslinked cyclodextrin polymers possess excellent passive resistance to protein, cell, and bacterial attachment, likely due to the hydrophilic and electrically neutral surface properties of these coatings. At the same time, anti-biofouling performance decreased with increasing crosslink ratios, possibly a reflection of decreased polymer mobility, increased rigidity, and increased hydrophobic character. Cyclodextrin-based materials may be broadly useful as coatings in industrial or medical applications where biofouling-resistant and/or drug-delivering surfaces are required.

## I. Introduction

Biological fouling, or “biofouling,” is the undesired accumulation of biological contaminants (biofoulants) on a material surface, particularly at an aqueous liquid/solid interface. Biofouling poses major challenges in the health care^1^, water treatment^2^, and marine industries^3^, among others^4,5^. For example, medical implants are susceptible to uncontrolled surface accumulation of proteins, cells, and bacterial biofilms, resulting in serious complications such as foreign body response^6,7^, thrombosis^8,9^, and prosthetic infection^1,10,11^. Water purification and desalination membranes are vulnerable to biofilm colonization that reduces flux and contaminates treated water^2,12,13^. Buildup of aquatic life on ship hulls accelerates corrosion, reduces vessel maneuverability/speed, raises fuel consumption, promotes invasive species migration, and necessitates periodic dry-docking maintenance^3,14–16^.

Biofouling is a cumulative process that often starts with non-specific protein adsorption^17–19^. Immediately upon exposure to a bare surface, protein solutes adsorb in an equilibrium determined by concentration, diffusivity, and affinity^17^. Small, abundant proteins predominate on the surface over short time frames, then are gradually replaced by higher-affinity proteins, a process known as the Vroman effect^20^. With few exceptions, proteins adsorb onto surfaces in a near-monolayer arrangement^17^. This “conditioning film” then mediates further biofouling events – typically cell adhesion, then matrix/biofilm formation as cells deposit new proteins, then macro-organism attachment (in the case of marine biofouling) – dependent on the types, concentrations, and conformations of the adsorbed molecules^15,21–24^. For this reason, surfaces that prevent protein adsorption are theorized to block cell adhesion^15,25^. Conversely, increased adsorption is typically expected to enhance cell adhesion^17,26^. However, many findings challenge this correlation^27,28^, indicating the importance of testing materials in biofouling scenarios that recapitulate conditions of intended use.

Anti-biofouling (ABF) materials possess surfaces that resist accumulation of proteins, cells, and/or organisms^25,29^, thus reducing consequences linked to excessive buildup of biological material on critical interfaces. ABF has typically been achieved by two major strategies: active ABF and passive ABF. These strategies are based on degrading adherent biofoulants or preventing their attachment, respectively^8,22^.

Active ABF approaches use biocidal agents presented at, or released from, the material surface to injure or destroy any cells or organisms that stick^30^. Typical agents presented at medical implant surfaces include silver compounds and antibiotic drugs. Biocides released from marine surfaces include toxic metal compounds^14,16,31–33^. Key limitations of active ABF strategies include: 1.) potential for off-target effects detrimental to surroundings, and 2.) transient effectiveness, given the finite biocide reservoir and the propensity for target micro-organisms to develop resistance. Additionally, depending on the biodiversity encountered, the biocidal agents chosen may not provide a sufficiently broad effect to resist biofouling by all relevant organisms.

Passive ABF approaches utilize anti-adhesive material surfaces to reduce the ability of biological contaminants to physically settle and adhere. Upon wetting, the foulability of a surface is influenced by numerous factors, including fluid movement near the interface, the area of surface exposed, the duration of biofoulant exposure, and the surface chemistry, charge, wettability, stiffness, and topography^13,14,23,34^. Surfaces that resist biofouling typically do so by minimizing contact (e.g. by trapping an air layer at the surface to minimize wetted area, or by attracting a water layer that sterically hinders protein adsorption) and attractive forces (e.g. electrostatic) between biofoulants and the surface.

The wettability of a surface plays a large role in the onset of protein adsorption. Specifically, hydrophobic materials have a strong tendency to adsorb and unfold (i.e. denature) protein solutes^13,25,35–39^. It is energetically favorable for proteins to displace polar water molecules at a hydrophobic surface, and further to change conformation such that core nonpolar amino acid segments are exposed and able to associate with the surface via nonpolar interactions. Even superhydrophobic substrates, which are hydrophobic materials that possess a nano-structured surface that traps air to minimize wetted area and biofoulant exposure, are prone to irreversible wetting and subsequent biofouling with extended submersion in protein solutions^16,40–43^. Beyond serving as a conditioning layer for further biofouling, denatured proteins on a hydrophobic medical device may elicit thrombotic, fibrotic, or inflammatory responses that limit the device’s biocompatibility^39,44–51^.

Conversely, hydrophilic material surfaces attract water molecules that sterically hinder protein adsorption while minimizing the chance of protein denaturation. Pioneering studies have indicated that electrically-neutral hydrophilic surfaces, especially those presenting abundant H-bond acceptors without H-bond donors, tend to be the most protein-resistant^52^. Overall charge neutrality minimizes electrostatic interactions that could otherwise attract proteins (or cells) to a surface^53,54^. The importance of lacking H-bond donors is less obvious^55^ as glycocalyx-mimetic carbohydrate surfaces (which display many H-bond donors) also effectively resist non-specific protein adsorption^56–58^. It is clear, however, that water held at the surface through H-bonding is critical for resistance to protein adsorption. For this reason, neutrally-charged, H-bond-acceptor-rich, hydrophilic polymers, such as polyzwitterions^59^, poly(ethylene glycol) (PEG)^60,61^, poly(2-hydroxyethyl methacrylate) (pHEMA)^62^, and many polysaccharides^56–58^ present key advantages for passive ABF. The major limitation of passive ABF strategies is that their performance may be compromised by surface defects (e.g. cracking or delamination of passive ABF coating exposes underlying substrate to biofoulants).

Our group has previously studied polymers composed of cyclic oligosaccharides, in which cyclodextrin (CD) molecules are crosslinked together to form insoluble polymer networks^63^. CDs are toroid-shaped molecules with a hydrophilic exterior and relatively hydrophobic core. They have long been used in the food and pharmaceutical industries, being Generally Recognized as Safe by the US FDA, to solubilize or prevent aggregation of nonpolar compounds in aqueous mixtures. CD polymers have principally been investigated for application as implantable drug delivery depots, based on the unique ability of CD subunits to reversibly bind and release drug compounds (e.g. antibiotics) through non-covalent (“affinity”) interactions^64^. These affinity properties make CD polymers very well-suited for active ABF, because when compared to purely diffusion-based active ABF materials, CD polymers can release biocides over longer durations^63,65,66^, and can be refilled more easily once the reservoir is depleted^67^. Additionally, given their neutrally-charged, H-bond acceptor-rich composition of hydrated saccharide units, CD polymers share many similarities with glycocalyx-mimetic passive ABF materials. Furthermore, proteins such as albumin have demonstrated weak affinity for cyclodextrin subunits^68^. These observations imply a possibility that CD polymers might be useful for passive ABF in addition to active ABF, a unique property not seen in traditional ABF materials^69^. This potential has not been sufficiently explored and could be of value for many medical and industrial applications. Therefore, the objective of this work was to assess protein adsorption, and attachment of mammalian and bacterial cells on CD polymer surfaces.

An overview of this work is presented in **Figure 1**. In this study, polymerized CD (pCD) was applied as a coating for polypropylene (PP), chosen as a model substrate material because of its susceptibility to uncontrolled biofouling in medical^70–74^ and industrial^75–80^ applications. Given the low surface energy of olefin polymers, PP substrates were treated with nonthermal plasma to enhance pCD coating uniformity and adherence^81^. We hypothesized that pCD coatings would deter protein adsorption, cell adhesion, and bacterial attachment to PP substrates, in a manner dependent on crosslinking. In this case, hexamethylene diisocyanate (HDI) was chosen as a crosslinker in order to allow investigation of neutrally-charged pCD materials having different overall hydrophobicity. Four different crosslinking formulations of pCD were first characterized in terms of material properties: swellability, rigidity, and wettability. Chemical composition of these formulations was also characterized using Fourier-transform infrared (FTIR) spectroscopy, and the impact of antibiotic drug loading on pCD material properties was preliminarily examined. Next, the four pCD formulations were studied in terms of their passive ABF performance relative to bare PP and polystyrene (PS) control surfaces in terms of protein adsorption from bovine plasma, mammalian fibroblast adhesion and viability, and bacterial (*S. aureus, E. coli*) attachment.

**Figure 1:**
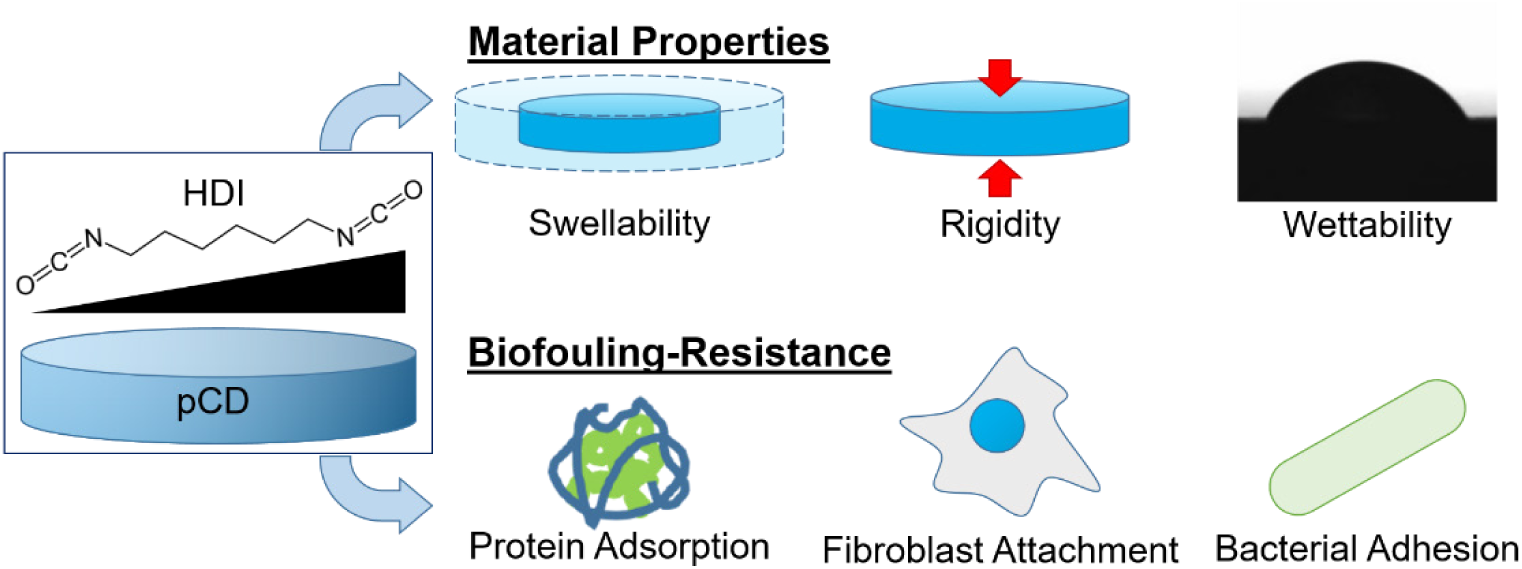
Study overview. The effects of HDI crosslinking on pCD material/surface properties (swellability, rigidity, wettability) and resistance to biofouling (protein adsorption, fibroblast attachment, bacterial adhesion) were explored.

## II. Methods

### 1. Materials

Soluble, lightly epichlorohydrin-crosslinked β-CD polymer precursor (bCD) was purchased from CycloLab R&D (#CY-2009, batch CYL-4160, MW ∼116 kDa; Budapest, Hungary). HDI crosslinker (#52649), black PP 96-well plates (#M9685), pHEMA (#P3932), and Pluronic F108 (#542342) were purchased from Sigma Aldrich. N,N-dimethylformamide (DMF) solvent (#D119-4), tissue culture PS (TCPS) 24-well plates (#08-772-1H), non-TCPS 12-well plates (#08-772-50), Dulbecco’s Modified Eagle Medium (DMEM) with high glucose (#11-995-040), fetal bovine serum (#16-141-079), penicillin-streptomycin (#15-140-122), Trypsin-EDTA (#25-200-056), LB broth (#BP1426-500), and 1.5 mL tubes (#05-408-129) were purchased from Fisher Scientific. BBL broth (#211768) and agar (#214010) was purchased from Becton Dickinson. PP 24-well plates (#1185U58) and lids (#1185U62) for fibroblast attachment were purchased from Thomas Scientific. Poly(tetrafluoroethylene) (PTFE) evaporating dishes with inner diameters 63 mm (#355314-0025) and 30 mm (#355304-0025) were obtained from Lab Depot. The antibiotic rifampicin (#R64000) was purchased from Research Products International. Sterile bovine plasma with sodium heparin anticoagulant (#IBV-N) was purchased from Innovative Research. PP sheet stock (#8742K133) for wettability and protein adsorption, and rod stock (#8658K51) for *S. aureus* attachment, were purchased from McMaster-Carr. Guava ViaCount reagent was purchased from Millipore Sigma (#4000-0040). PP 4-0 Prolene blue suture (#8592G) was purchased from eSutures. Bioluminescent ilux pGEX(-) *E. coli* was purchased from Addgene, a gift from Dr. Stefan Hell (plasmid #107879)^82^. Methicillin-resistant *S. aureus* strain Xen30 was purchased from Caliper Life Sciences. Tissue homogenizer (#TH-01) and blades (#30750H) were purchased from Omni International.

### 2. Plasma cleaning and activation of PP substrate surfaces

To maximize pCD coating adherence and stability, PP substrates were placed in a 4” diameter x 8” length quartz reaction chamber of a Branson/IPC Model #1005-248 Gas Plasma Cleaner and treated with nonthermal plasma (500 mTorr, 50 W, 13.56 MHz) using an inlet gas mixture of argon bubbled through water (Ar/H2O)^81,83^. PP substrates were treated for a fixed 10 min duration within 1 h of pCD coating application or direct biofoulant exposure (for plasma-treated bare PP controls). Non-treated (0 min) PP samples without any known prior exposure to plasma or ultraviolet light were included as controls in all experiments.

### 3. pCD synthesis and coating onto surfaces

In order to examine the impact of crosslinking on material properties and ABF performance, pCD was synthesized using HDI as a crosslinker for bCD at approximate crosslinker / glucose residue molar ratios of 0.08, 0.16, 0.32, and 0.64 (Table 1). bCD was weighed and placed in PP tubes, then DMF was added to dissolve it at 33% w/v. HDI was added to solutions to achieve the desired crosslink ratios, and pre-polymer mixtures were thoroughly vortexed, then cast either: (i) into clean PTFE dishes to produce free films of pCD for subsequent punching of disks for measurement of swelling ratio, elastic modulus, and contact angle, (ii) onto flat PP sheet/rod stock pieces (newly abraded using 1200, 2500, then 5000 grit SiC sandpaper to expose fresh surface) for preparing coated specimens for protein adsorption and *S. aureus* attachment, (iii) into wells of PP multiwell plates for production of coated well surfaces for measurement of mammalian fibroblast adhesion and viability. For coated well surfaces, the volume of pre-polymer mixture added was 140 µL/well for 24-well plates, and 42 µL/well for 96-well plates, then plates were agitated to promote complete coverage, and surfaces were visually examined post-curing to exclude defective coatings prior to use. Cast pre-polymer mixtures were kept covered with Parafilm and typically allowed to cure for at least 4 days at ambient temperature and pressure. Cured pCD coatings were rinsed several times to terminate crosslinking, and stored immersed in sterile PBS (for subsequent cell/bacterial culture) or deionized water (for XPS) to keep samples hydrated before use.

**Table 1:** Pre-polymer mixtures for different formulations of pCD*. * HDI volumes (density 1.047 g/mL) are calculated based on estimation that epichlorohydrin linkages and water comprise a negligible weight fraction of bCD.

| pCD Formulation | bCD (mg) | DMF (μL) | HDI (μL) |
| --- | --- | --- | --- |
| **0.08** | 1000 | 3000 | 80 |
| **0.16** | 1000 | 3000 | 160 |
| **0.32** | 1000 | 3000 | 320 |
| **0.64** | 1000 | 3000 | 640 |

## 4. Effects of HDI-crosslinking on pCD physicochemical properties

### 4.1. Determination of pCD swelling ratio

Crosslinking density impacts the ability of network polymers to absorb water, and the degree of swelling may influence the biofouling resistance of ABF polymers^84^. For this reason, effects of HDI crosslinking on pCD swelling ratio were examined. pCD pre-polymer mixtures (scaled to a DMF volume of 1 mL) were cast into 30 mm diameter PTFE evaporating dishes, covered, and allowed to cure for 7 weeks at ambient temperature and pressure. Disks were punched out 6.0 mm in diameter using a biopsy punch, extracted from dishes, and hydrated in deionized water for 3 days. Hydrated disks (n = 10-12 per group) were gently blotted on filter paper to remove surface water, and weighed after 60 seconds to the nearest 0.1 mg (wet weight) on an analytical balance (Mettler Toledo ME104TE). Samples were then placed onto sheets of Parafilm and dried overnight under vacuum at ambient temperature, and weighed again to the nearest 0.1 mg (dry weight) on the same balance. Swelling ratio for each individual disk was calculated as the absorbed water weight (wet weight minus dry weight) divided by the dry weight. Results shown represent findings from one experiment, with the same trends having also been observed in 4 similar independent experiments.

### 4.2. Unconfined compression testing of pCD

Crosslinking directly alters network polymer rigidity, and biomaterial stiffness is recognized as a factor that impacts adhesion, spreading, and phenotype of mammalian cells^85,86^. Therefore, effects of HDI crosslinking on pCD mechanical properties were examined. pCD pre-polymer mixtures (scaled to a DMF volume of 4.5 mL) were cast into pristine 63 mm diameter PTFE evaporating dishes, covered, and allowed to cure for either 4 days or 4 weeks at ambient temperature and pressure. Disks were then punched out 8.0 mm in diameter using a biopsy punch (1.38-2.26 mm thickness, n = 5-9 per group) and placed in deionized water, with which specimens were kept hydrated before and throughout testing. For preparation of drug-loaded specimens, disks were incubated for 24 h in 5 mL DMF containing 5% w/v rifampicin (RIF), then rinsed numerous times and stored for >24 h in beakers of deionized water. Specimen dimensions were carefully measured with digital calipers to the nearest 0.01 mm before each test and crosshead speed was adjusted accordingly to ensure a consistent strain rate of −0.005 s^-1^. Unconfined compression tests were performed on a Rheometrics RSA II Solids Analyzer (Rheometric Scientific) up to a load limit of 10 N (50 kPa) using a data acquisition rate of 4 Hz. Elastic modulus was determined through linear regression of the stress-strain curve over all possible 2.5% strain ranges within each given test prior to reaching the load limit (or the onset of yielding for some pCD 0.08 specimens), with the maximum slope found being defined as the modulus. Results shown represent pooled findings from 2 independent experiments.

### 4.3. Contact angle goniometry

Because HDI crosslinks in pCD can be expected to be nonpolar and wettability is known to affect biofouling resistance, the effect of HDI crosslinking on pCD wettability was examined. pCD pre-polymer mixtures (scaled to a DMF volume of 4.5 mL) were cast into pristine 63 mm diameter PTFE evaporating dishes, covered, and allowed to cure for 4 days at ambient temperature and pressure. Disks were punched out 12.0 mm in diameter. Specimens were kept hydrated with deionized water until contact angle measurement. For preparation of drug-loaded specimens, disks were incubated for 24 h in 5 mL DMF containing 5% w/v RIF, then rinsed numerous times and stored for >24 h in beakers of deionized water. Surfaces were evaluated for wettability by static contact angle measurement using a KSV Instruments CAM 200 Optical Contact Angle Meter. Sample surfaces were gently blotted to remove surface moisture on dry filter paper followed by KimWipes, and then deionized water droplets of 8 µL volume were dispensed onto each sample surface and allowed to equilibrate for 30 s prior to photographing and measurement. Two unique droplets per sample were measured for 3 separate samples per group, and the measurement for each droplet reflects the average of the angles on the left and right sides. Measurements were performed using KSV CAM 2008 software. Results shown represent findings from one experiment, with the same trends having also been observed in 2 similar independent experiments.

### 4.4. Attenuated Total Reflectance FTIR Spectroscopy

FTIR was performed on pCD surfaces to confirm the extent to which reactants are consumed in polymerization and verify that HDI crosslink ratio is reflected in polymer composition. After cured pCD disks were punched and extracted from PTFE dishes, the polymer film remnants were re-covered, allowed to cure at ambient temperature until 4 weeks had elapsed since casting, then removed from the dishes and dried for 2 days under vacuum. pCD films were characterized using an Excalibur FTS 3000 FTIR spectrometer (BioRad, Hercules, CA) equipped with a Pike MIRacle single-reflection attenuated total reflectance (ATR) accessory, germanium crystal, and flat-tipped pressure anvil (Pike Technologies). Scans were collected (4 cm^-1^ resolution, 800-4000 cm^-1^ range, 5 kHz speed, sensitivity 16, open aperture, 100 co-added scans, and Boxcar apodization function) on 3 samples per group while dry nitrogen gas was continuously flowed to purge the system of CO2 and water vapor. Background scans were collected before the first sample, and absorbance values relative to background were converted to % transmittance. In Microsoft Excel, spectra were averaged to include all samples within each group, shifted to set the average value between 1900-2200 cm^-1^ to 100%, and normalized such that the magnitude of the 1037 cm^- 1^ peak attributed to stretching of the CD ether (C-O-C) functionality^87,88^ was held constant across all CD-based groups to account for differences in crystal-sample contact. The curve for HDI was normalized such that the peak at 2926 cm^-1^ corresponding to alkane C-H stretch was equal in magnitude to that in the curve for .64 pCD.

## 5. Effects of HDI-crosslinking on pCD anti-biofouling performance

### 5.1. Evaluation of protein adsorption

Given the important role of protein adsorption in subsequent biofouling processes, we attempted to characterize the resistance of pCD to protein adsorption. Protein adsorption was assessed using subtractive X-ray Photoelectron Spectroscopy (XPS) for the following surfaces: bare TCPS, bare untreated PP, and bare plasma-treated PP (included as controls), and plasma-treated PP surfaces coated with 0.08, 0.16, 0.32, and 0.64 pCD. Two samples were prepared per group. PP sheet stock was cut to dimensions of ∼7.5 mm x 7.5 mm, gently sanded to expose fresh surface using a graded series (1200, 2500, then 5000 grit) of SiC sandpaper, and rinsed thoroughly with deionized water. Plasma treatments and pCD coatings were then applied to the appropriate samples. After curing of pCD, all samples were hydrated in deionized water. One sample per group was then incubated in a sterile undiluted solution of heparinated bovine plasma for 24 h at 37°C on a rotisserie shaker, while the other sample per group remained in a bath of deionized water. After incubation, protein-exposed samples were then washed thoroughly with deionized water. All samples were evaluated for elemental content using a PHI Versaprobe 5000 Scanning X-Ray Photoelectron Spectrometer equipped with Al Kα source (hν = 1486.6 eV). Survey scans were collected using 200 µm spot size, 45 W power, 15 kV acceleration voltage, 117.40 eV pass energy, 0.40 eV step size, 25 ms/step, 8 cycles, 44.7° take-off angle, and 0-1100 eV range. The C1s peak was auto-shifted to 284.8 eV, and the ratios of carbon, nitrogen, and oxygen were analyzed. Peak areas were taken with background set using a Shirley function from 280-292 eV for C1s, 396-404 eV for N1s, and 526-538 eV for O1s. Analysis was performed using MultiPak software version 9.8.0.19 (Physical Electronics, Inc.). The amount of nitrogen detected was considered indicative of both the amount of adsorbed protein, as well as the degree of HDI-crosslinking of pCD polymers on coated samples. Therefore, protein adsorption onto surfaces was measured indirectly based on the difference between the percentage of N/(N+C+O) after incubation in bovine plasma versus without incubation in bovine plasma. Results shown represent findings from one experiment.

### 5.2. Investigation of mammalian cell attachment and viability

Attachment of host cells to an implant surface determines the extent of tissue ingrowth around the biomaterial, and it is critical that the material be safe for these cells so as to promote a favorable host response. Attachment of mammalian cells and viability of attached cells were assessed using hemacytometry and flow cytometry for the following surfaces: bare TCPS, bare PS, bare untreated PP, and bare plasma-treated PP (included as controls), and plasma-treated PP surfaces coated with 0.08, 0.16, 0.32, and 0.64 pCD. NIH/3T3 fibroblasts were suspended in sterile DMEM with 10% fetal bovine serum and 1% penicillin-streptomycin, and seeded onto culture surfaces at a supra-confluent density of 150,000 cells/cm^2^. Cells were allowed to attach for either 3 h or 21 h at 37°C prior to rinsing twice with 750 µL sterile media, then once with 750 µL sterile PBS. Cells were then enzymatically detached from surfaces using 200 µL 0.25% Trypsin-EDTA followed by 2 further rinses of 500 µL sterile media. Rinsates and trypsinates were retained in separate tubes and analyzed using flow cytometry on a Millipore Guava EasyCyte using Guava ViaCount as a viability indicator according to manufacturer instructions. Results shown represent pooled findings from 5 independent experiments for 3 h hemacytometry, and 2 independent experiments each for 21 h hemacytometry, 3 h viability, and 21 h viability (1 sample per experiment).

### 5.3. Measurement of *S. aureus* attachment

Attachment of bacteria to a biomaterial surface is a key step in prosthetic infection, therefore we investigated the ability of both gram-positive (*S. aureus*) and gram-negative (*E. coli*) bacteria to adhere to pCD. *S. aureus* attachment was assessed using colony-forming unit (CFU) counts. This was done for bare untreated PP control, and plasma-treated PP coated with 0.16 pCD. This was the only pCD formulation tested in this experiment given the low-throughput nature of CFU analysis, and was chosen for its demonstrated ABF performance in mammalian cell attachment and protein adsorption experiments. PP rod stock with a ¼” diameter was cut into cylinders of lengths between 2-3.5 mm (n = 4-5 per group). Cut cylindrical samples were sanded smooth (to ensure comparable surface topography between samples) using a graded series of SiC sandpaper. The length and diameter of each individual sample were recorded to the nearest 0.01 mm using digital calipers. Samples were submerged in 70% ethanol and allowed to dry, then plasma treatments and coatings were applied to the appropriate samples. After coatings had been allowed to cure, all samples were hydrated in sterile deionized water. Xen30 *S. aureus* bacteria were thawed from frozen stock, inoculated into a 14 mL round-bottom tube of sterile BBL broth, and expanded in suspension for 24 h in a dedicated 37°C incubator while the tube lid was vented. The tube was then removed from the incubator and stored at 4°C to maintain bacteria in the stationary phase. Prior to seeding onto experimental surfaces, the bacterial suspension was diluted to an OD600 of 0.72 relative to sterile BBL broth. The bacterial concentration at this OD600 is estimated as 2×10^12^ mL^-1^ based on CFU counts of dilutions spread onto agar plates. The bacterial suspension was then directly seeded at 1 mL per sample onto non-coated and coated cylinders in 1.5 mL tubes, then incubated at 37°C for 24 h on a rotisserie shaker. Following incubation, cylinders were rinsed by dipping in 5 sequential 2 mL baths of sterile PBS and then suspended in 1 mL sterile BBL broth and ground at 30,000 rpm with a tissue homogenizer using separate sterile blades for each individual sample. The bacterial concentration in each homogenized suspension was then quantified by spreading dilutions onto surfaces of agar-coated plates and performing CFU counts after 18 h at 37°C. The number of adherent bacteria per square mm of sample surface area was then calculated. Results shown represent findings from one experiment, with the same trend having also been observed in one similar independent experiment.

### 5.4. Assessment of *E. coli* attachment

*E. coli* attachment was assessed using bioluminescence measurements for the following 96-well plate surfaces: bare untreated PP, bare plasma-treated PP, bare TCPS, untreated PP coated with Pluronic F108, pHEMA-coated untreated PP, and plasma-treated PP coated with 0.08, 0.16, 0.32, and 0.64 pCD. The additional controls Pluronic F108 and pHEMA were included solely in this experiment because bacterial attachment is involved in the majority of biofouling challenges encountered in both medicine and industry, and because bioluminescence measurement is high-throughput, thus facilitating their study. All coatings (pCD pre-polymer mixtures, sterile-filtered 2% w/v pHEMA in 95% ethanol, and sterile-filtered 2% w/v Pluronic F108 in PBS) were applied at 42 µL/well. pHEMA coatings were left uncovered in a sterile hood and allowed to dry overnight before being covered with Parafilm and stored. Pluronic coatings were covered with Parafilm for 2 days, then rinsed several times and stored immersed in sterile PBS. Bioluminescent ilux pGEX(-) *E. coli* were thawed from frozen stock, inoculated into a 14 mL round-bottom tube of sterile lysogeny broth (LB), and expanded in suspension for 24 h in a dedicated 37°C incubator with the tube lid vented. The tube was then removed from the incubator and stored at 4°C to maintain bacteria in the stationary phase. Prior to seeding onto experimental surfaces, the bacterial suspension was diluted to an optical density at 600 nm (OD600) of 0.50 relative to sterile LB broth. The bacterial concentration at this OD600 is estimated as 10^11^ mL^-1^ based on CFU counts of dilutions spread onto agar plates. The bacterial suspension was then directly seeded at 100 µL/well, cultured under static conditions for 24 h at 37°C, and following incubation, wells were rinsed 5 times with 200 µL/well sterile LB broth, then emptied and filled a final time with 100 µL/well sterile LB broth. Bioluminescence measurement was then performed using a Biotek Synergy H1 plate reader (∼23°C read temperature, luminescence endpoint scan, 5 s integration time, 1 mm read height, full light emission, 135 gain, top optics, 100 ms delay, extended dynamic range). A subset of non-rinsed wells for each type of surface were also seeded in duplicate with several known concentrations of bacteria, included for creation of standard curves. Results shown represent pooled findings from 2 independent experiments (n = 16-32 total wells per group).

## 6. Statistical Analysis

All data is presented as mean ± standard deviation. Statistical significance was defined for all analyses as p<0.05. Bivariate correlations were evaluated using Spearman’s rho (rS) in Minitab 2019. All other data comparisons were performed using 2-tailed 2-sample t-tests with unequal variance in Microsoft Excel 2016.

## III. Results

### 1. Effects of HDI-crosslinking on physicochemical properties of pCD

Given that the overarching goal of this work was to explore the usefulness of pCD for ABF applications, we first sought to broadly characterize the material properties of several different CD polymer formulations that might potentially be applied towards ABF purposes. The effects of crosslinking on swelling ratio, elastic modulus, contact angle, and chemical composition of pCD were thus explored. Additionally, considering that these polymers might even be used for simultaneous passive ABF and active ABF or drug delivery, we also assessed effects of drug-loading on elastic modulus and contact angle of pCD. To this end, the antibiotic rifampicin (RIF), was chosen as a model drug, due to its poor water-solubility, and known ability to form inclusion complexes with CD subunits.

Swelling ratio describes the ability of network polymers to absorb water, and the degree of swelling may reflect polymer chain mobility, therefore influencing ABF performance^84^. The swelling ratio of CD polymers was found to decrease with increasing HDI crosslinking ratio (rS = −0.968, p < 0.001) (**Fig 2a**). This could be a result of the decreasing mobility of the polymer network upon further crosslinking.

**Figure 2:**
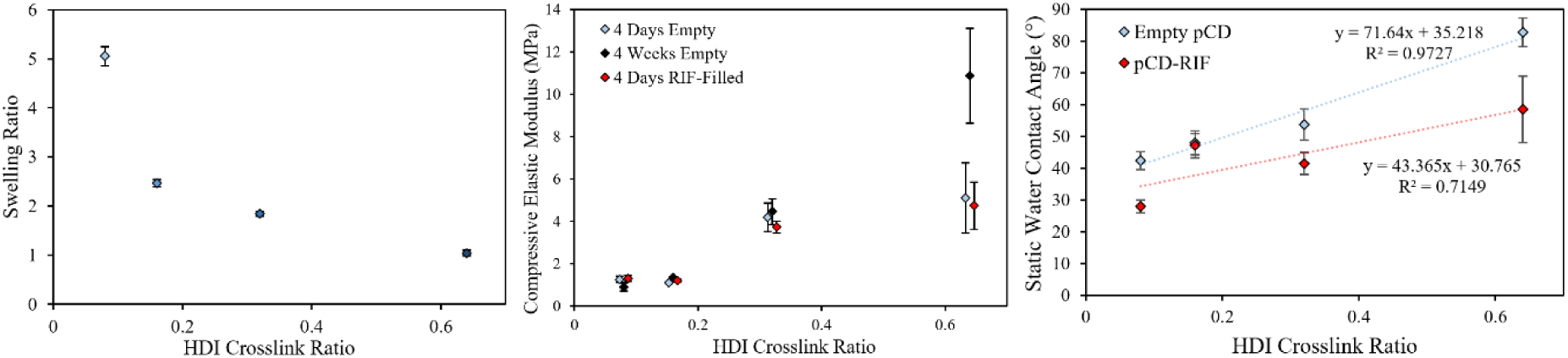
Physical properties of 0.08, 0.16, 0.32, and 0.64 pCD formulations: (a) swelling ratio, (b) elastic modulus, (c) static water contact angle. Points for elastic modulus and contact angle are intentionally staggered along the abscissa to improve readability.

Biomaterial stiffness (i.e. elastic modulus) is considered to be a factor which impacts adhesion and spreading of cells. In agreement with findings of decreasing swelling ratio, elastic moduli of CD polymers were found to increase with increasing HDI crosslinking ratio (**Fig 2b**). This was true for empty pCD after 4 days (rS = 0.802, p < 0.001) or 4 weeks (rS = 0.969, p < 0.001) of crosslinking, and for RIF-filled pCD after 4 days of crosslinking (rS = 0.796, p < 0.001). Increasing the duration of crosslinking from 4 days to 4 weeks led to a doubling in elastic modulus at the 0.64 crosslink ratio (p < 0.001), reflecting increased formation of covalent crosslinks within the pCD network over this time frame. Drug-loading of RIF had no meaningful effect on elastic modulus at any crosslink ratio (p > 0.07). All pCD samples except for 43% of those in the 4-day 0.08 pCD empty group, and 13% of those in the 4-day 0.08 pCD-RIF group, were able to sustain loads up to the 10N load limit without apparent yielding or failure. This indicates possible fragility of the polymers at the lowest crosslinking ratio. Incorporation of RIF into pCD was previously shown to increase compressive properties of pCD microparticle-laden bone cement composites on the order of MPa, an effect thought to be in part due to RIF complexation hindering structural collapse of molecular pockets within pCD^89^. Results here, however, suggest that RIF has little direct effect on the compressive moduli of isolated pCD materials.

Surface wettability is recognized as an important property that affects biofouling resistance of a material. Static water contact angles of CD polymers increased with increasing HDI crosslink ratio (**Fig 2c**), likely as a consequence of incorporation of nonpolar hexamethylene spacers with further crosslinking. This was apparent for both empty (rS = 0.825, p < 0.001) and RIF-filled pCD (rS = 0.473, p < 0.017). This indicates increasing hydrophobicity of pCD with further incorporation of nonpolar HDI crosslinks. Interestingly, incorporation of the poorly water-soluble RIF into pCD led to decreased static water contact angles at most crosslink ratios (p < 0.015) except for 0.16 (p = 0.753). This result may reflect diffusion of RIF into the water droplet, or increased presentation of polar functionalities that conceal nonpolar HDI crosslinks at the surface as RIF molecules complex with CD subunits.

Normalized FTIR spectra (**Fig 3**) revealed that increased crosslinking ratio led to relatively larger peak sizes at 1256cm^-1^, 2856 cm^-1^, and 2926 cm^-1^ corresponding to alkane C-H stretch^87,88,90– 92^, at 3332 cm^-1^ corresponding to both hydrogen-bonded alcohol O-H and secondary amide N-H stretch^90,93^, at 1576 cm^-1^ corresponding to urethane N-H stretch^87,94^, at 1624 cm^-1^ corresponding to urethane C=O bending^87^, at 1278 cm^-1^ corresponding to urethane C-N stretch^90^, and at 1460 cm^-1^ corresponding to C-H bending^92^. Together, these peak size increases confirm increased abundance of HDI crosslinks with increased crosslink ratio. Additionally, for pCD spectra, disappearance of the peak seen in the HDI spectrum at 2270 cm^-1^ indicates that isocyanates were fully converted (most likely to urethanes) in crosslinking reactions^90,91,94,95^.

**Figure 3:**
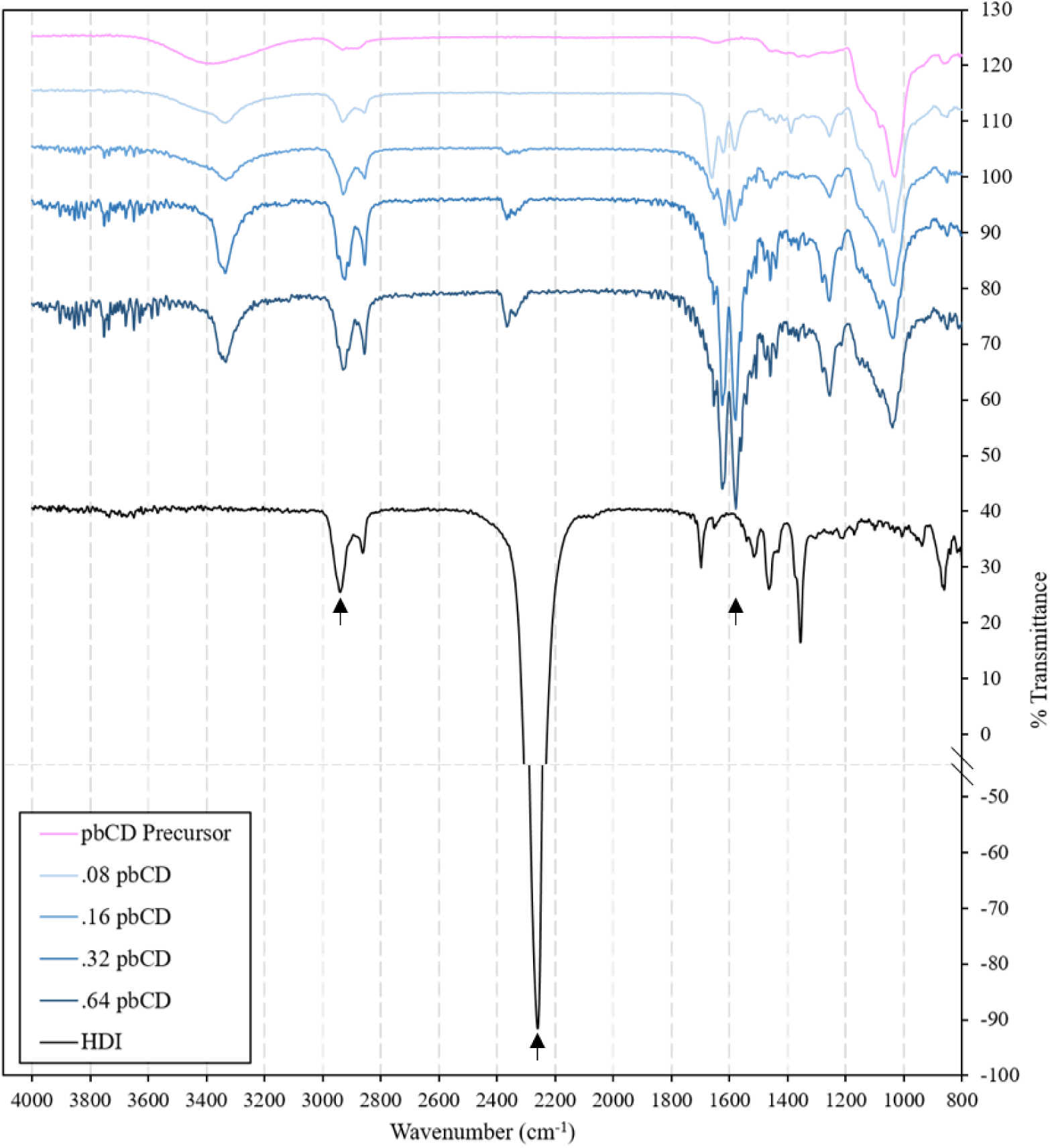
Normalized ATR-FTIR spectra of pCD materials and components. Spectra are intentionally shifted along the ordinate to improve readability. Arrows are drawn at 2926 cm^-1^ for the alkane C-H stretch peak, 2270 cm^-1^ for the isocyanate peak only seen in the HDI spectrum, and 1576 cm^-1^ for the urethane N-H stretch peak which appears only after pCD crosslinking.

### 2. Effects of HDI-crosslinking on pCD resistance to non-specific protein adsorption

Having characterized the effects of HDI-crosslinking on the physical properties of pCD, we next sought to evaluate protein adsorption on the surfaces of these polymers. Subtractive XPS was used to measure protein adsorption onto polymer surfaces. This method was chosen over colorimetric or fluorimetric protein binding assays because several of our preliminary studies indicated that the tags and labels (e.g. FITC, Coomassie brilliant blue) used to visualize proteins in such experiments had inherent affinity for CD, thus making such strategies insufficiently specific and sensitive for detection of protein adsorption (data not shown). Prior literature corroborates these observations^96,97^.

XPS measurements suggested that protein adsorption onto pCD polymers was lower than that onto TCPS or PP (**Fig 4**). Protein adsorption onto pCD was observed to decrease with decreasing crosslink ratio, possibly as a result of increasing hydrophilic character. The coating for the 0.08 pCD surface was found to have delaminated during a rinse step, so no data is shown for that sample. Plasma treatments were observed to reduce protein adsorption onto PP surfaces, in agreement with previous findings^83^, again possibly a result of increasing wettability.

**Figure 4:**
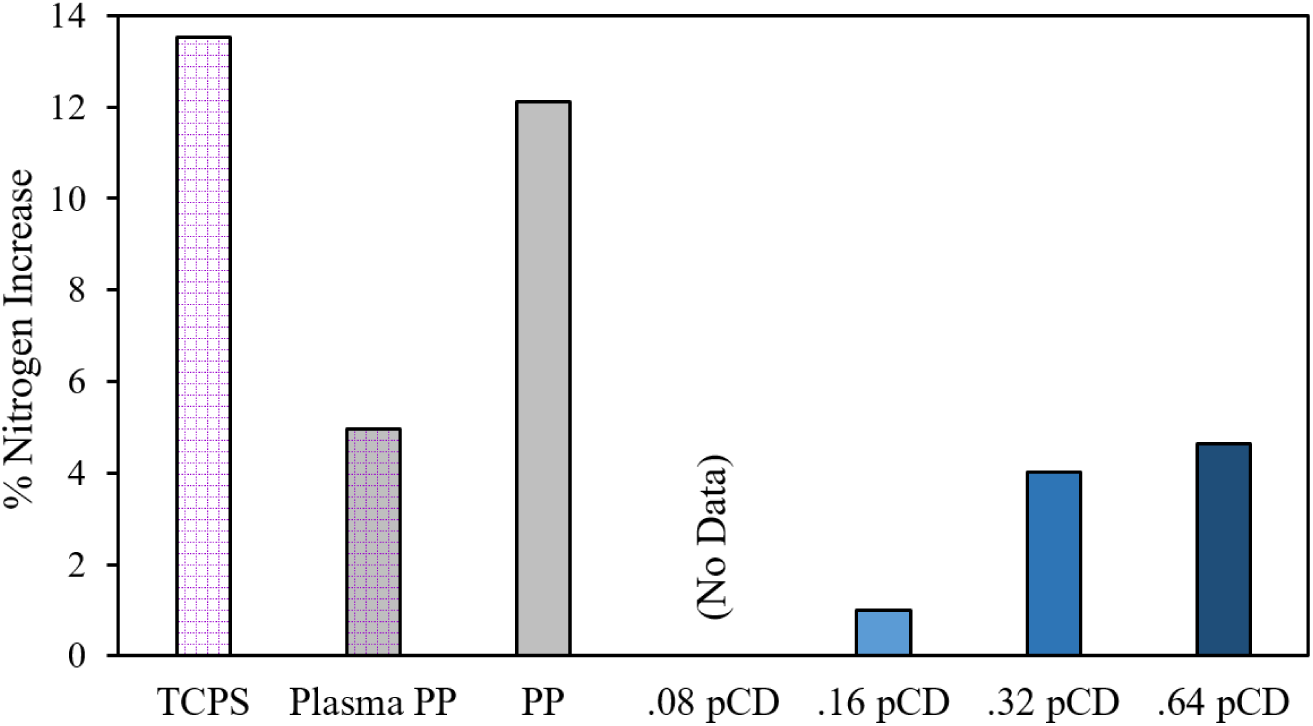
Protein adsorption onto surfaces as measured using subtractive XPS. Increased nitrogen content reflects increased protein adsorption.

### 3. Effects of HDI-crosslinking on pCD resistance to mammalian cell adhesion

Following protein adsorption experiments, mammalian cell (NIH/3T3 fibroblast) adhesion and viability were studied using hemacytometry and flow cytometry, respectively. These methods were chosen over optical or colorimetric *in situ* measurements because our preliminary studies suggested that indicator dyes for cell metabolism (e.g. Alamar blue) had inherent affinity for CD, thus interfering with assay results (data not shown). Prior literature corroborates this observation^98^. Furthermore, while certain CD-based polymers can be made optically transparent^99^, the translucent PP substrates and CD polyurethane coatings in this study were not optimized for cell observation.

Cell adhesion for pCD polymers was found to range from 0-20% of the level of cell adhesion observed for TCPS (p < 0.001) (**Fig 5a**). Among pCD polymers, the most crosslinked, 0.64 formulation was found to possess the least resistance to cell adhesion (p < 0.003). This indicates that lightly-crosslinked pCD polymers resisted stable adhesion of fibroblasts, likely due to low levels of protein adsorption. Cell adhesion was equally low for untreated PP as for most pCD polymers (p > 0.293). Plasma treatment of PP increased cell adhesion (p = 0.01), consistent with prior findings^83^. Likewise, cell adhesion for TCPS (commercially plasma-treated PS) was higher than that for untreated PS (p < 0.003). The low cell adhesion to untreated PP may indicate that the identities and/or conformations of the serum proteins which adsorb onto the PP surface are unfavorable for cell adhesion. Additionally, despite relatively lower apparent levels of protein adsorption on plasma-treated PP, the adsorbed proteins on this material seem to be more favorable for cell adhesion than those on untreated PP. The high standard deviations for plasma PP and untreated PS appeared to a result of the fibroblasts forming monolayers and either detaching all at once, or remaining attached altogether, resulting in an apparent bimodal distribution for cell adhesion in these groups.

**Figure 5:**
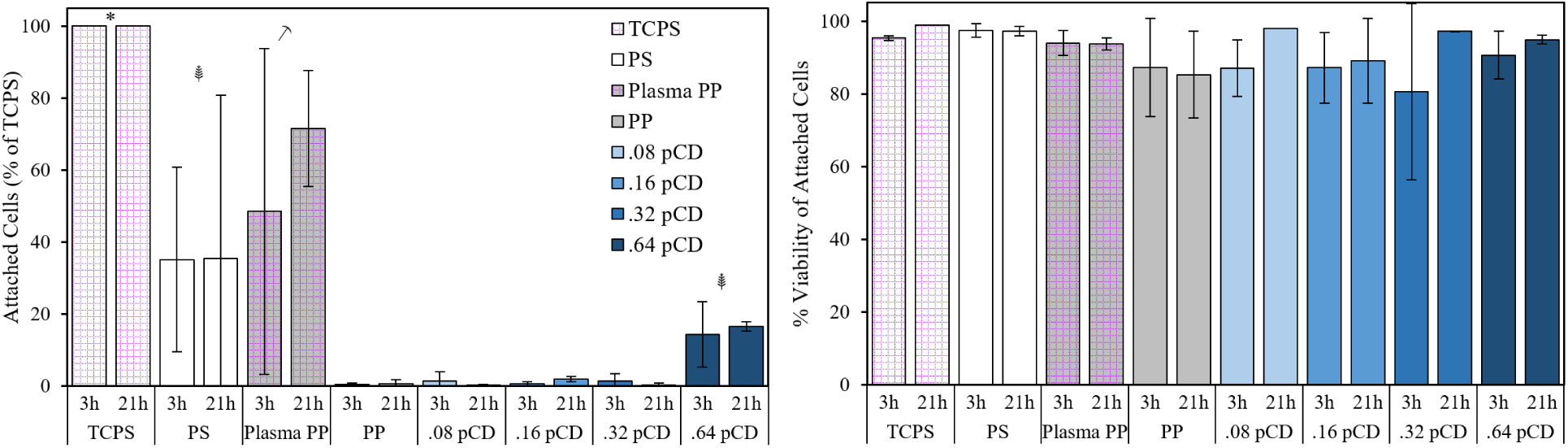
Cell counts (a), and viability (b), following 3h or 21h of adhesion to control and pCD surfaces. Data reflect only the cells that attached to pCD, so as to exclude effects of anoikis-induced cell death. *Denotes significant difference to all other groups. 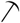 Denotes significant difference to all pCD groups and untreated PP. 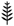 Denotes significant difference to untreated PP and .08, .16, and .32 pCD. No differences were observed between groups in terms of viability.

Viability of the cells that did attach to pCD polymers was comparable to that of controls (p > 0.08) (**Fig 5b**), with all values >75%, suggesting that pCD polymers did not exert cytotoxic effects on cells. This result is encouraging, as previous studies have indicated that at high concentrations, non-polymeric CD can reduce cell viability by abstracting cholesterol from cell membranes^100^. Polymerization of CD monomers may help mitigate such cytotoxic effects. Similar trends for cell counts and viability were observed at both 3h and 21h.

### 4. Effects of HDI-crosslinking on pCD resistance to bacterial cell attachment

Having evaluated protein adsorption and mammalian cell adhesion onto pCD, bacterial attachment was next investigated. This was done in two separate sets of experiments, the first for the gram-positive species *S. aureus*, in this case the methicillin-resistant Xen30 strain, and the second for the gram-negative species *E. coli*, specifically the bioluminescent ilux pGEX(-) strain.

*S. aureus* attachment to .16 pCD after 24h was significantly lower than attachment to untreated PP (p = 0.041) (**Fig 6a**). Likewise, *E. coli* attachment to plasma-treated PP, .08 pCD, .16 pCD, .32 pCD, and PP coated with Pluronic F108 after 24h was significantly lower than attachment to untreated PP (p < 0.001) (**Fig 6b**). *E. coli* attachment to surfaces coated with Pluronic F108, chosen a control because of its ease of application and highly effective passive ABF properties^101^, was roughly half that seen for .08 and .16 pCD (p < 0.014). Surprisingly, pHEMA did not resist attachment of *E. coli* as well as TCPS or PP (p < 0.001), despite its common use for creating coatings on cultureware that prevent adhesion of mammalian cells. This could be a reflection of differences in the mechanisms by which *E. coli* and anchorage-dependent mammalian cells adhere to surfaces. In support of this explanation, plasma treatment suppressed attachment of *E. coli* despite enhancing attachment of NIH/3T3 fibroblasts.

**Figure 6:**
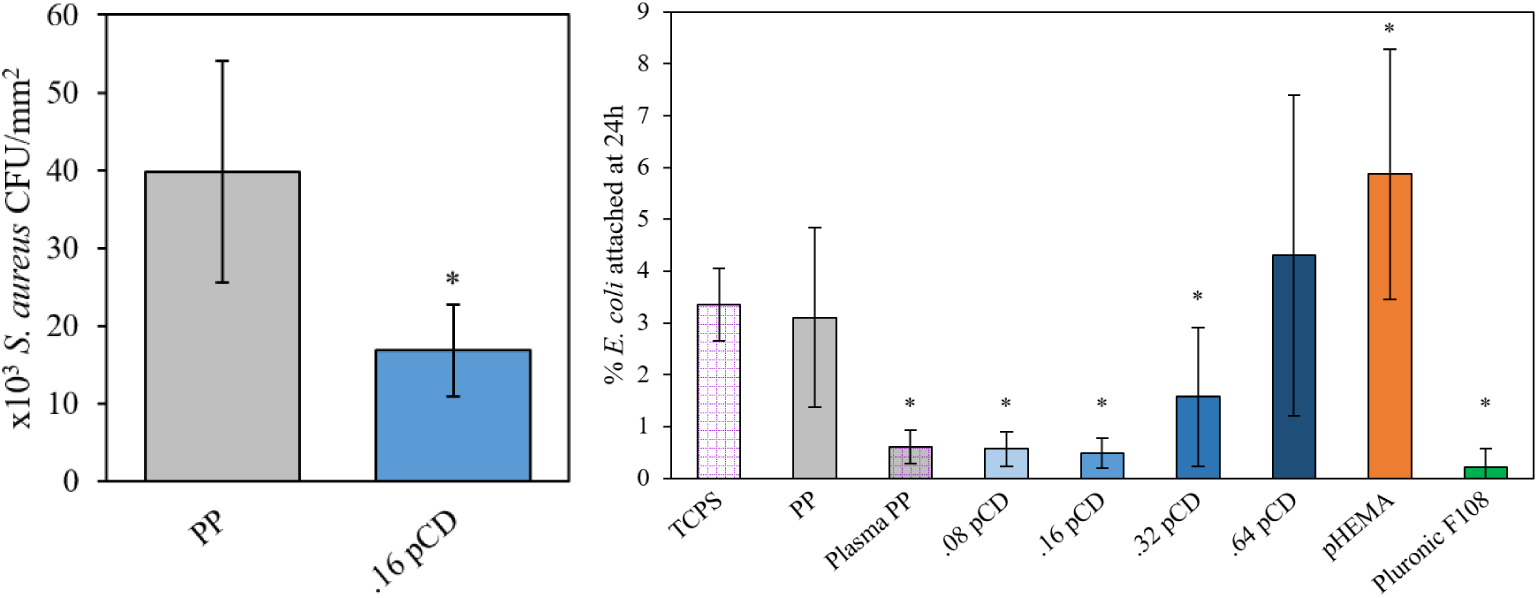
Bacterial attachment of *S Aureus* (a) and *E coli* (b). *Represents significant difference (p<0.05) to PP.

## IV. Discussion

This study has demonstrated that CD polymers, in addition to their ability to actively release biocidal compounds, have potential to passively resist biofouling by proteins, mammalian cells, and bacteria, without apparent cytotoxicity. Increased HDI crosslinking of CD polymers was found to decrease swellability, increase rigidity, and increase hydrophobicity, but at the same time tended to reduce pCD resistance to protein, cell, and bacterial attachment. This reduction in ABF properties with increased HDI crosslinking might be expected given that less wettable materials have a stronger tendency to promote protein adsorption^13,25,35–39^ (along with subsequent biofouling events), that higher substrate rigidity promotes mammalian cell adhesion and spreading^85,86^, and that more abundant crosslinks should restrict chain mobility and limit presentation of polar domains at the network polymer surface. Although this study design did not allow for independent determination of the degree to which each material property individually affected biofouling resistance of pCD, results here provide insights into the rational design of pCD for ABF purposes and set the stage for further exploration.

Carbohydrate surface coatings, such as dextran, cellulose, or mannose, have previously been investigated for ABF applications^56–58^, but unlike CD, these sugars lack the ability to form affinity-based inclusions of small molecule drugs. Affinity properties give pCD materials unique advantages for active ABF, such as more sustained drug release^63,65,66^ that is relatively independent of material dimensions^102^, and more efficient biocide refilling^67^. PEG-based coatings may be the most popular materials studied for ABF purposes. Results from *E. coli* repulsion experiments suggest that lightly-crosslinked pCD formulations may perform nearly as well in terms of resistance to bacterial adhesion as a freshly PEGylated surface. However, PEG can be prone to oxidation in the presence of oxygen and metal ions found in physiological solutions^103^, and many human patients have shown antibodies to PEG following its increasing use in food, cosmetic, and pharmaceutical products^104^, potentially limiting its utility for long-term or medical ABF applications. Cyclodextrins, conversely, do not elicit immune responses in mammals^64^, and the CD subunits and urethane bonds in pCD are in theory quite stable. Zwitterionic polymers are another class of polymers that have become popular for ABF research given outstanding performance, their main drawback being that they are difficult and costly to synthesize^53^. pCD on the other hand is easy to make, and inexpensive.

Apart from ABF, a unique advantage of pCD is its ability to complex with and deliver small-molecule drug cargo over long periods of time. It may seem counterintuitive that a polymer having inherent affinity for small-molecule drugs could be useful for prevention of protein adsorption, cell adhesion, and bacterial attachment. One possible explanation is the much smaller size of drug compounds – which must be able to fit in a molecular pocket having a diameter of 5.7Å, 7.8Å, or 9.5Å for the native cyclodextrins α-, β-, and γ-CD, respectively^105^ – than proteins, bacteria, and cells. Most proteins, which are typically at least one order of magnitude larger than a small-molecule drug (yet still much smaller than bacteria or cells), are likely too bulky to diffuse through the pCD polymer matrix let alone complex with CD subunits. Additionally, unlike small-molecule drugs, proteins in aqueous solution exist in conformations that inherently shield their nonpolar domains against hydrophobic interactions with CD subunits. Similarly, polyurethane materials such as pCD undergo surface restructuring in aqueous solution through microphase separation, which enriches the polymer surface with hydrophilic domains^106^, further deterring protein displacement of water and adsorption. With increased HDI crosslinking, the proportion of hydrophobic segments in pCD increases, while their mobility (i.e. ability to be shielded from protein solutes at the interface) decreases, with the consequence being increased protein adsorption and subsequent biofouling.

Interestingly, incorporation of the poorly water-soluble antibiotic RIF into pCD increased wettability and had minimal effect on material rigidity. Considering that increased wettability might generally be expected to improve passive ABF properties, these results together may suggest that drug loading with biocidal agents, like RIF, for purposes of active ABF does not negatively influence the passive ABF properties of pCD.

Nonthermal plasma treatments of PP in this study, just as in a prior study^83^, were shown to reduce non-specific protein adsorption and *E. coli* attachment, while enhancing adhesion of mammalian fibroblasts, all of which could be favorable outcomes for a plasma-treated biomedical implant. However, the effects of plasma treatment are known to diminish (i.e. age) over time. Literature suggests that hydrophobic recovery of plasma-treated PP normally occurs over the course of several weeks, the rate being dependent on environmental factors and polymer crystallinity^107–110^. The presence of a coating may act to protect or constrain the activated PP substrate surface.

One limitation of this study is that the effects of different crosslinkers or subunits were not examined. A more hydrophilic crosslinker might in theory produce polymers that remain resistant to biofouling regardless of crosslink ratio. In this study, HDI was chosen as a crosslinker to allow investigation of neutrally-charged polymers having different overall hydrophobicity. In preliminary studies, a different neutrally-charged crosslinker, ethylene glycol diglycidyl ether, was found to produce polymers that swelled to the point of rupture and delamination upon hydration post-curing. The polymers produced by HDI-crosslinking were more robust, therefore we considered HDI a more useful crosslinker for preparation of pCD coating materials here. There are many other possible crosslinkers that could be considered^90,111–114^. Similarly, different CD subunits were not evaluated in this study. Thus, we were not able to comprehensively assess the generality of the passive ABF performance of pCD. β-CD was considered the subunit of choice among the native cyclodextrins given that from a drug delivery standpoint, it is the most widely used and generally has the most versatility in terms of the variety of guest molecules that it can complex with^115–118^. However, each of the native cyclodextrins possess different dimensions and water solubilities, thus changing the subunit could foreseeably alter polymer affinity properties and hydrophilicity, as well as ABF performance. For instance, Dos Santos *et al*. showed that incorporation of β- and γ-CD decreased lysozyme and albumin adsorption onto pHEMA hydrogels, while incorporation of α-CD decreased lysozyme adsorption but increased albumin adsorption^119^.

We envision that pCD might be applied as coatings for medical devices, such as textile implants. Specifically, CD-based polymer coatings have previously been investigated for use as surgical mesh coatings that could enable controlled antibiotic/drug release^65,87,120–123^, but it would also be noteworthy to explore the potential of these materials to prevent post-surgical adhesions. The formation of tissue adhesions on a surgical fabric surface is preceded by two events that CD-based polymers are shown to be capable of mitigating: 1.) non-specific adsorption and denaturation of protein (especially fibrinogen), which promotes coagulation, inflammation, and impaired fibrinolysis, and 2.) adhesion of fibroblasts that remodel persistent coagulation products into fibrous connections between the implant surface and adjacent tissue structures. Alternatively, pCD coatings may be useful for application onto PP water treatment membranes. Such coatings might simultaneously resist bacterial biofouling and remove small-molecule pollutants from water^124^. This investigation provides a basis for the use of CD-based polymers for such applications. Future studies should seek to directly evaluate field or *in vivo* performance of pCD coatings for ABF purposes.

## V. Summary and Conclusions

This study has demonstrated across several different readouts the potential of cyclodextrin-based polymers to passively resist biofouling by proteins, mammalian cells, and bacteria, without overt cytotoxicity. Additionally, the effects of HDI crosslinking on pCD swellability, rigidity, and wettability were characterized. Lower HDI-crosslinking densities yielded pCD with superior resistance to biofouling, an effect that could be attributed to increased chain mobility, decreased rigidity, and/or decreased hydrophobicity. Cyclodextrin-based coating materials may be appropriate in industrial or medical applications for which biofouling-resistant and/or drug-delivering surfaces are required.

## VI. Acknowledgments

The authors gratefully acknowledge support from National Institutes of Health: NIH R01GM121477 (HvR), and NIH NIAMS Ruth L. Kirschstein NRSA T32 AR007505 Training Program in Musculoskeletal Research (GDL). Core facility services provided by the Swagelok Center for Surface Analysis of Materials, and the Cytometry & Microscopy core at Case Western Reserve University are also appreciated. The authors also thank Joseph Mansour for use of the Rheometrics instrument, the Advincula group for use of the contact angle goniometer, Kevin Abbassi for expertise and assistance with XPS, David Jasen WuWong for assistance with *S. aureus* bacterial attachment experiments, Katherine Yan for assistance with sample preparation, and Erika Cyphert and Ali Ansari for writing suggestions.

